# Establishment and Application of a duplex real-time qPCR method for detection of *Salmonella* spp. and *Serratia fonticola* in imported feedstuffs

**DOI:** 10.1101/324798

**Authors:** Jing hua Ruan, Wu jun Wang, Quan yang Bai, Ti yin Zhang, Teng zheng, Shi yu Yu, Zhi deng Zhang, Su jie Lin, Bo xiong Zhong, Dao jin Yu

**Affiliations:** College of Animal Science, Zhejiang University, Hangzhou, P. R. China; Fujian Key Laboratory of Traditional Chinese Veterinary Medicine and Animal Health, Fujian Agriculture and Forestry University, Fuzhou, P. R. China; Fujian Key Laboratory for Technology Research of Inspection and Quarantine, Fujian Entry-exit Inspection and Quarantine Bureau, Fuzhou, P. R. China

**Keywords:** *Salmonella* spp., *S. fonticola*, Duplex real-time qPCR, Imported feedstuffs, Feed safety, Bacterial pathogens

## Abstract

*Salmonella* spp. is a high-risk bacterial pathogen that is monitored in imported animal-derived feedstuffs. *Serratia fonticola* is the bacterial species most frequently confused with *Salmonella* spp. in traditional identification methods based on biochemical characteristics, which are time-consuming and labor-intensive, and thus unsuitable for daily inspection and quarantine work. In this study, we established a duplex real-time qPCR method with *invA*-and *gyrB*-specific primers and probes corresponding to *Salmonella* spp. and *S. fonticola*. The method could simultaneously detect both pathogens in imported feedstuffs, with a minimum limit of detection for *Salmonella* spp. and *S. fonticola* of 197 copies/μL and 145 copies/μL, respectively (correlation coefficient R^2^ = 0.999 in both cases). The amplification efficiency for *Salmonella* spp. and *S. fonticola* was 98.346% and 96.49%, respectively. Detection of clinical samples was consistent with method GB/T 13091-2002, and all 20 artificially contaminated imported feed samples were positively identified. Thus, the developed duplex real-time qPCR assay displays high specificity and sensitivity, and can be used for the rapid and accurate detection of genomic DNA from *Salmonella* spp. and *S. fonticola* within hours. This represents a significant improvement in the efficiency of detection of both pathogens in imported feedstuffs.

**Importance:** Imported feedstuffs must be tested for pathogenic *Salmonella* species that represent a biological hazard. Various *non-Salmonella* colony-forming species belong to *Enterobacteriaceae,* and *Serratia fonticola* forms colonies of similar color and morphology to *Salmonella* spp., leading to confusion in daily quarantine tests. Traditional methods based on biochemical and serological characteristics are cumbersome and labor-intensive, and unable to fully support current quarantine testing demands. Thus, there is an urgent need to develop a rapid and accurate method for the effective identification of these pathogens. The duplex real-time qPCR method established herein can rapidly identify *Salmonella* spp. and *S. fonticola*, and has great potential for application to feed safety and prevention of exterior pathogens.

## Introduction

*Salmonella* spp. are ubiquitous Gram-negative bacteria in the environment and include six different subspecies and more than 2000 serotypes that infect a wide range of hosts, often causing severe food poisoning outbreaks in humans and other animals. People infected with *Salmonella* can develop diarrhea, fever, and suffer dehydration, hence *Salmonella* spp. are of significance to public health.

*Serratia fonticola* is a species belonging to the *Serratia* genus that was first isolated from water and soil in 1979 (1). Subsequent studies showed that *S. fonticola*, a member the Gram-negative *Enterobacteriaceae* family that includes *Salmonella* spp., is also ubiquitous in environments such as water, soil, plants, and the gastrointestinal tract of humans and other animals (2). Research has revealed that *S. fonticola* can infect various tissues and organs in humans (3, 4), causing septic arthritis (5), septicemia (6, 7), gastrointestinal tract infections (8, 9), and surgical infections (10). Therefore *S. fonticola* have been defined as an important opportunistic pathogen.

At present, the detection of *Salmonella* spp. in imported animal-derived feeds (fish meal and chicken powder) involves non-selective enrichment, selective enrichment, selective platelet culturing, biochemical culturing of suspected *Salmonella* colonies (triglyceride tests, etc.), and even serological typing. These conventional methods require at least 3 days to detect *Salmonella* spp. in daily quarantine work. Furthermore, suspected *Salmonella* colonies based on selective plate isolation and culturing of imported animal-derived feeds often turn out to be S. *fonticola* when biochemical properties are investigated (11). The enzyme linked immune sorbent assay (ELISA) method is commonly used for detecting pathogens in imported animal-derived feeds, but this method can produce incorrect results due to its cumbersome operational steps. A novel method involving polymer fluorescent nanoparticles as biosensor probes has been described for the detection of *Salmonella* (12), which takes only 3 h, but cannot be applied to large volume detection due to its prohibitively high cost.

PCR is widely used in the detection of pathogens due to its high sensitivity and specificity, and results can often be obtained within several hours. Numerous traditional PCR and multiplex PCR methods have been developed for the detection of species belonging to both the *Salmonella* and *Serratia* genera (13–17). Multi-PCR approaches can differentiate two or more pathogens in one amplification, however, products are easily contaminated during agarose gel electrophoresis, which increases false-positive and false-negative results. Thus, a more reliable approach such as real-time fluorescence quantitative PCR (RT-qPCR) would be more desirable for the rapid and accurate detection of pathogens. RT-qPCR methods are known to be fast, reliable, and highly efficient. Several single RT-qPCR methods for detecting *S. nematodiphila* (18), *S. marcescens* (19–21), and *Salmonella* spp. (22–24) have been described, and multiplex RT-qPCR detection methods have also been reported (25–28).

Previous studies on fish meal and other animal-derived feeds have mainly focused on the detection and epidemiology of *Salmonella*, *Shigella*, *Escherichia coli*, and other common pathogenic bacteria. A method for the simultaneous detection of *Salmonella* spp. and *S. fonticola* in imported animal-derived feeds using RT-qPCR has not been reported. Such a method could provide rapid differential diagnosis of suspected *Salmonella* colonies after selective plate separation and culturing. This would undoubtedly improve the detection of *Salmonella*, and greatly shorten the time required for subsequent biochemical and serological identification, saving valuable manpower and material resources.

The duplex real-time qPCR method established in the present work provides a useful tool for the simultaneous detection of *Salmonella* spp. and *S. fonticola.* The method has important theoretical significance and great potential for improving the safety of imported feeds by rapidly identifying bacterial pathogens and facilitating effective quarantining in a more timely manner than traditional detection methods.

## Materials and Methods

### Bacterial strains

A total of 48 tested strains were used in this study, including 17 reference strains from six different collection centers, and 31 isolates from imported fishmeal (Table 1, 2, and 3). All experimental strains were streaked on nutrient agar plates and cultured in Luria-Bertani (LB) broth at 37°C overnight (~18-24 h), except for *S. marcescens,* which was grown at 23°C for 24 h. An established single colony was inoculated into 3 mL of LB broth for 8 h and the resultant culture was harvested for genomic DNA extraction.

**TABLE 1.** *Salmonella* spp. strains used in this study

| Bacterial species | Strain No. | No. of strains | Source of strains |
| --- | --- | --- | --- |
| *Salmonella enteritidis* | ATCC13076 | 1 | ATCC (Manassas, USA) |
| *Salmonella typhimurium* | GDM1.237 | 1 | GDMCC (Guangdong, China) |
| *Salmonella montevideo* | / | 2 | Imported fishmeal |
| *Salmonella weltevreden* | / | 1 | Imported fishmeal |
| *Salmonella enteritidis* | / | 3 | Imported fishmeal |
| *Salmonella birkenhead* | / | 1 | Imported fishmeal |
| *salmonella anatis* | / | 1 | Imported fishmeal |
| *Salmonella senftenberg* | / | 1 | Imported fishmeal |
| *Salmonella lomita* | / | 1 | Imported fishmeal |
| *Salmonella saintpaul* | / | 1 | Imported fishmeal |
| *Salmonella* O:2 | / | 3 | Imported fishmeal |
| *Salmonella* O:4 | / | 1 | Imported fishmeal |
| *Salmonella* O:7 | / | 2 | Imported fishmeal |
| *Salmonella* O:8 | / | 2 | Imported fishmeal |
| *Salmonella* O:9 | / | 3 | Imported fishmeal |
| *Salmonella* O:3,10 | / | 5 | Imported fishmeal |

**TABLE 2.** *Serratia* genus strains used in this study

| Bacterial species | Strain No. | No. of strains | Source of strains |
| --- | --- | --- | --- |
| *Serratia fonticola* | GDM1.995 | 1 | GDMCC (Guangdong, China) |
| *Serratia marcescens* | MCCC1A06806 | 1 | MCCC (Xiamen, China) |
| *Serratia proteamaculans* | MCCC1K00532 | 1 | MCCC (Xiamen, China) |
| *Serratia odorifera* | GDM1.864 | 1 | GDMCC (Guangdong, China) |
| *Serratia ficaria* | GDM1.994 | 1 | GDMCC (Guangdong, China) |
| *Serratia plymuthica* | GDM1.996 | 1 | GDMCC (Guangdong, China) |
| *Serratia rubidaea* | GDM1.1008 | 1 | GDMCC (Guangdong, China) |
| *Serratia liquefaciens* | CICC21538 | 1 | CICC (Beijing, China) |
| *Serratia grimesii* | ACCC01695 | 1 | ACCC (Beijing, China) |
| *Serratia fonticola* | / | 3 | Imported fishmeal |

**TABLE 3.**
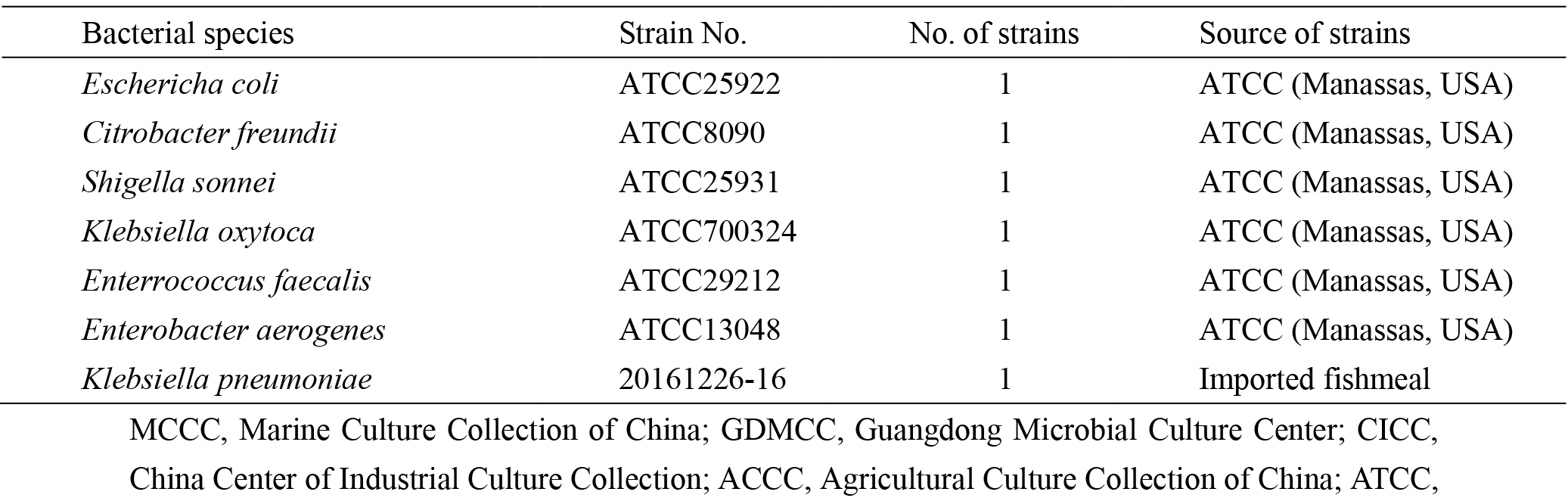
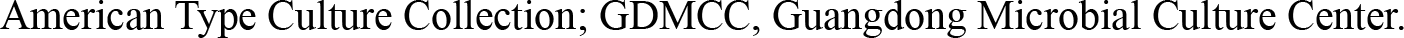
Other strains used in this study

| Bacterial species | Strain No. | No. of strains | Source of strains |
| --- | --- | --- | --- |
| *Escherichia coli* | ATCC25922 | 1 | ATCC (Manassas, USA) |
| *Citrobacter freundii* | ATCC8090 | 1 | ATCC (Manassas, USA) |
| *Shigella sonnei* | ATCC25931 | 1 | ATCC (Manassas, USA) |
| *Klebsiella oxytoca* | ATCC700324 | 1 | ATCC (Manassas, USA) |
| *Enterococcus faecalis* | ATCC29212 | 1 | ATCC (Manassas, USA) |
| *Enterobacter aerogenes* | ATCC13048 | 1 | ATCC (Manassas, USA) |
| *Klebsiella pneumoniae* | 20161226-16 | 1 | Imported fishmeal |

MCCC, Marine Culture Collection of China; GDMCC, Guangdong Microbial Culture Center; CICC,
China Center of Industrial Culture Collection; ACCC, Agricultural Culture Collection of China; ATCC,
American Type Culture Collection; GDMCC, Guangdong Microbial Culture Center.

### Sample collection and bacterial isolation

A 25 g sample of fishmeal was added to 225 mL of buffered peptone and incubated at 36 ± 1°C for between 16-20 h. A 10 mL sample of this pre-enrichment culture was transferred into 100 mL of enrichment solution containing selenite cystine, cultured at 36 ± 1°C for 24 h, and inoculated onto selective medium designed for *Salmonella* spp. Two *Salmonella* spp. colonies were cultured at 36 ± 1°C for 48 h in selective medium, one on the medium slope, the other punctured through the agar, and both were then inoculated on trisaccharide iron medium. Meanwhile, suspicious colonies were inoculated onto lysine decarboxylase medium and cultured at 36 ± 1°C for 18-24 h (or up to 48 h if necessary). Colonies from positive samples presumed to be *Salmonella* spp. were further identified using a VITEK II Compact 30 instrument (bioMérieux Technologies Inc., SA, French) with a GN card according to the manufacturer’s specifications.

### Species-specific primer and probe design

The *invA* sequences of *Salmonella* spp. were aligned to identify conserved and specific regions using CLUSTAL W software (29). A series of sense and antisense primers were designed based on these conserved and specific regions using ABI ViiA7 PrimerExpress software (Life Technologies Inc., Foster City, CA, USA), and the final specific primer pair and dual-labelled probe (Table 4) targeting the *Salmonella* spp. virulence gene were determined using the basic logical alignment search tool (BLAST) (30). Primers and dual-labelled probe targeting the *gyrB* gene were as described previously (31). Two primers and probe sets were synthesized by Sangon Biotech (Sangon Biotech Co., Ltd. Shanghai, China), and probes labeled with the fluorescent reporter dye carboxy-4′,5′-dichloro-2′,7′-dimethoxyfluorescein (JOE) targeting *invA* and 6-carboxyfluorescein (FAM) targeting *gyrB* were covalently coupled to the 5′-end, with Black Hole Quencher 1 (BHQ-1) at the 3′-end.

**TABLE 4.**
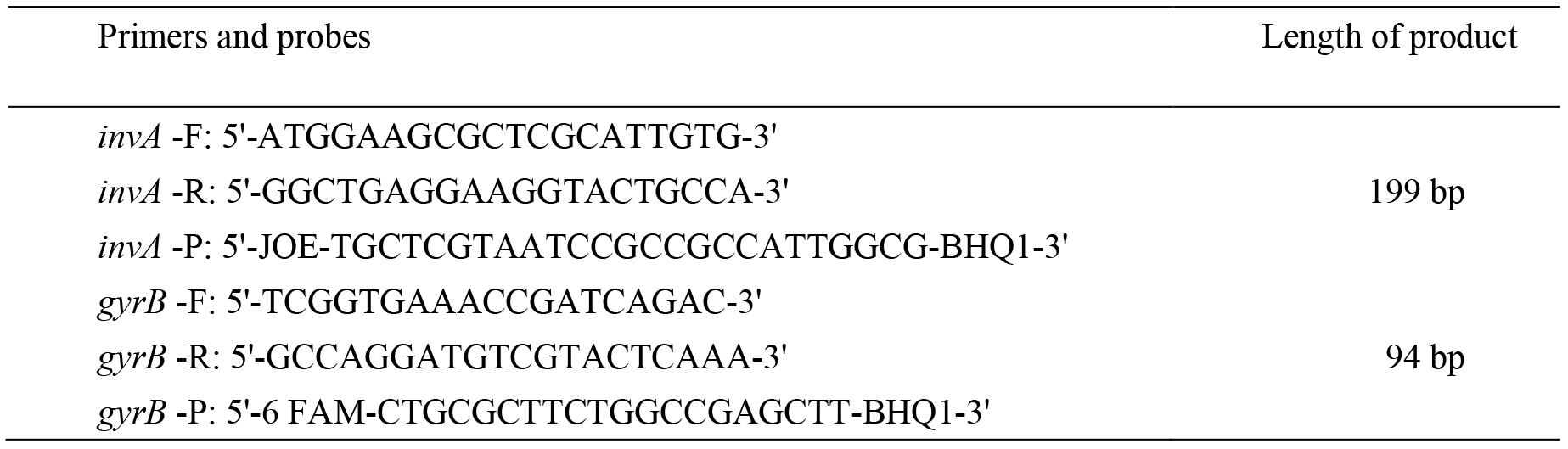
Real-time qPCR primer pairs and probes used in this study

### Extraction of bacterial genomic DNA

DNA extraction from isolates and reference strains was performed with a TIANamp Bacteria DNA Kit (Tiangen Biotech, Co., Ltd, Beijing, China) according to the manufacturer’s recommendations. DNA was eluted with sterile water and stored at −20°C until use. DNA purity and concentration were determined using a Quawell Q6000+ spectrophotometer (Quawell Technology, Inc., USA).

### Screening of the optimum annealing temperature of primers

The 20 μL qPCRs contained 10 μL of (2×) Premix Ex Taq (TaKaRa Biomedical Technology Co., Ltd, Beijing, China), 0.4 μL of each primer (10 μM) and probe (10 μM), 1 μL of bacterial DNA, and 7.8 of Rnase-free ddH_2_O. Reactions were performed on a Bio-rad CFX96 Real-time PCR system (Bio-Rad Laboratories, Inc., USA) using a two-step method. Thermal cycling conditions consisted of an initial denaturation at 95°C for 30 s, followed by 40 cycles at 95°C for 5 s and 65.0, 64.5, 63.3, 61.4, 59.0, 57.0, 55.7, or 55.0°C (triplicates for each temperature) for 34 s to determine the optimal annealing temperature of the two primer sets. CFX Manager Software (Bio-Rad Laboratories, Inc., USA) was employed to monitor PCR amplification, and the collect and analyze amplification data.

### Single real-time qPCR assays

Two simplex real-time qPCR detection methods for the detection of *Salmonella* spp. and *S. fonticola* were established using primers and probes targeting *invA* and *gyrB* genes, respectively, using an Applied Biosystems ViiA 7 real-time PCR system (Life Technologies Inc., Foster City, CA, USA). Reaction conditions were determined after various PCR parameters were tested according to the information supplied with the reagents. The optimized 20 μL PCR contained 10 μL of 2× Premix Ex Taq, 0.4 μL of each primer (10 μM) and probe (10 μM), 0.2 of ROX Reference Dye E (50×), 1 μL of bacterial DNA, and 7.6 μL of Rnase-free ddH_2_O. Optimized thermal cycling conditions involved an initial denaturation at 95°C for 30 s, followed by 40 cycles at 95°C for 5 s and 64°C for 34 s. ViiA 7 Software (Life Technologies Inc., Foster City, CA, USA) was employed to monitor PCR amplification, and the collect and analyze amplification data.

### Specificity of simplex real-time qPCR assays

To verify the specificity of the two simplex real-time qPCR assays, tests were performed by amplifying genomic DNA extracted from strains listed in Table 1, 2, and 3. Genomic DNA was extracted from 29 *Salmonella* spp. and 19 *non-Salmonella* species and used to determine the specificity of the designed primers and probes targeting the *invA* gene. The non-*S. fonticola* control containing 44 non-*S. fonticola* organisms, comprising eight different species belonging to the genus *Serratia*, and 36 *Enterobacteriaceae* strains, was used in RT-qPCR to confirm cross-reaction of *gyrB* gene-specific PCR primers and probes.

### Construction of recombinant plasmids

PCR products amplified with primers targeting *invA* and *gyrB* genes were purified using a DNA Fragment Purification Kit (Tiangen Biotech, Co., Ltd) after agarose gel electrophoresis, and sequenced by Sangon Biotech to confirm the identity of the target genes used in this study. The pMD19-T vector (Sangon Biotech) was used for cloning target genomic sequences, and the resultant constructs were transformed into competent *Escherichia coli* DH5α cells. Recombinant plasmids carrying each target gene were confirmed, and subsequent DNA sequencing was also performed by Sangon Biotech. Amplified sequences were compared with original sequences deposited in the GenBank database using NCBI BLAST.

### Sensitivity of single real-time qPCR assays

The purity and concentration of recombined plasmids pMD-invA and pMD-*gyrB* were determined using a Quawell Q6000+ spectrophotometer. Plasmids were 10-fold serially diluted nine times and subjected to real-time qPCR to construct standard curves (three technical replicates for each dilution), from which we determined both the amplification efficiency and the minimum detection limit of the two real-time qPCR methods. The plasmid copy numbers was calculated using the following formula (32): 
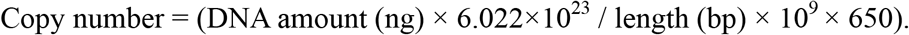

### Duplex real-time qPCR assay

A duplex real-time qPCR assay was established based on the two simplex RT-qPCR assays developed for the detection of *Salmonella* spp. and *S. fonticola* using an equal amount of genomic DNA from the two pathogens in the same reaction. Optimal PCR conditions were ultimately determined based on the simplex RT-qPCR parameters described above by varying a single factor while all other parameters remained constant. The main factor to be investigated was the concentration of the two primer and probe sets since the annealing temperatures of primers and probes was already calculated.

### Reproducibility and stability tests

To analyze the reproducibility and stability of the established duplex real-time qPCR method, recombinant pMD-invA and pMD-*gyrB* plasmids were 10-fold serially diluted five times (each dilution was tested in triplicate). A total of 96 reactions were simultaneously performed using the *invA-* and *gyrB* recombinant plasmids in the same real-time qPCR mixture to verify reproducibility. The coefficient of variation (CV) based on quantification cycle (Cq) values for each test was used to evaluate the performance of this approach.

### Detection of artificially contaminated imported feedstuffs

We tested 20 imported fishmeal samples not contaminated by either *Salmonella* spp. or *S. fonticola*. A 25 g portion of 10 of the fishmeal samples was artificially contaminated with *Salmonella* spp., while the other 10 samples were contaminated with *S. fonticola*, and contaminated samples were added to 225 mL of buffered peptone enrichment solution and incubated at 36 ± 1°C for 16-20 h. A 1 mL sample of the pre-enrichment culture was transferred into 10 mL of enrichment solution containing selenite cystine, cultured at 36 ± 1°C for 24 h, and 1 mL of the resultant culture was used for genomic DNA extraction using the TIANamp Bacteria DNA Kit (Tiangen Biotech, Co., Ltd). Finally, RT-qPCR was performed for the detection of *Salmonella* spp. and *S. fonticola,* and the national standard method (GB/T13091-2002) was also employed for verification.

### Detection of clinical imported feed samples

Ten imported fishmeal samples infected with *Salmonella* spp. and four imported fishmeal samples infected with *S. fonticola* were collected during quarantine work between June 2015 and June 2017. Genomic DNA was extracted from each sample after treatment as descripted above, real-time qPCR was performed for detection of these two pathogens, and the national standard method (GB/T13091-2002) was also employed for verification.

## Results

### Determination of optimum annealing temperature

Genomic DNA from *Salmonella enteritidis* and *S. fonticola* was used as a template to determine optimal annealing temperatures for the two primer sets by testing between 55 and 65°C using a Bio-Rad fluorescence quantitative PCR instrument. RT-qPCR using the designed primers and probes yielded the highest fluorescence intensity of amplified products at am annealing temperature of 64.5°C for both *invA-* and *gyrB*-specific primers and probes, and this annealing temperature was employed in subsequent experiments.

### Specificity of single real-time qPCR assays

The specificity of the two single real-time qPCR assays was verified by amplifying genomic DNA extracted from reference strains and laboratory isolates. As shown in Figure 1, the real-time qPCR assay detected *Salmonella* spp. only, while no fluorescent signal was observed for any *non-Salmonella* spp. strains or blank controls. However, the real-time qPCR assay targeting *S. fonticola* generated four specific amplification curves for the detection of four *S. fonticola* strains, while non-*S. fonticola* and blank controls were not amplified (Figure 2). Thus, the specificity of the two real-time qPCR assays was 100%, with no detectable fluorescent signal for negative samples or blank controls.

**FIGURE 1.**
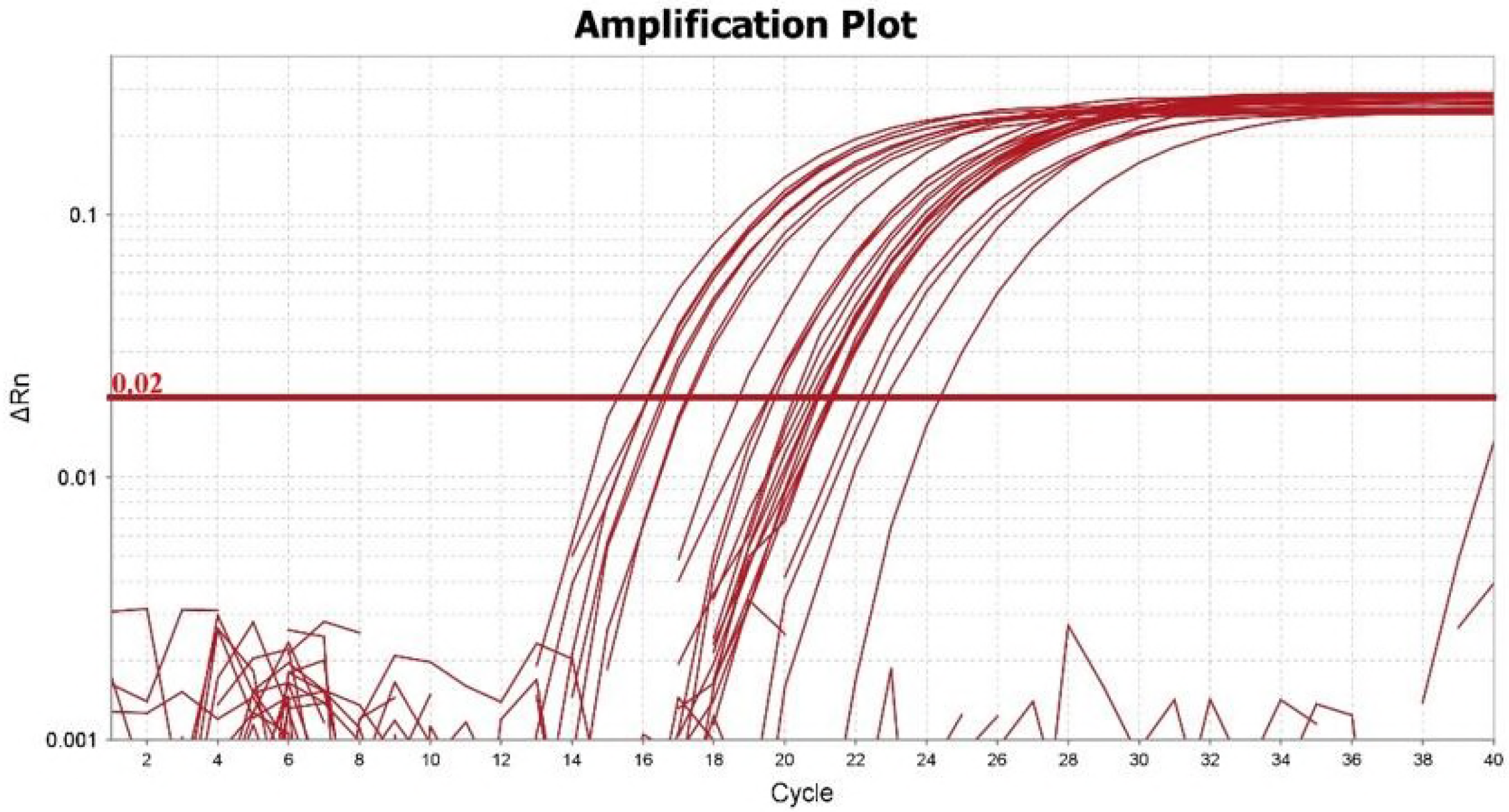
Specificity testing of the real-time qPCR assay using 29 *Salmonella* spp. strains.

**FIGURE 2.**
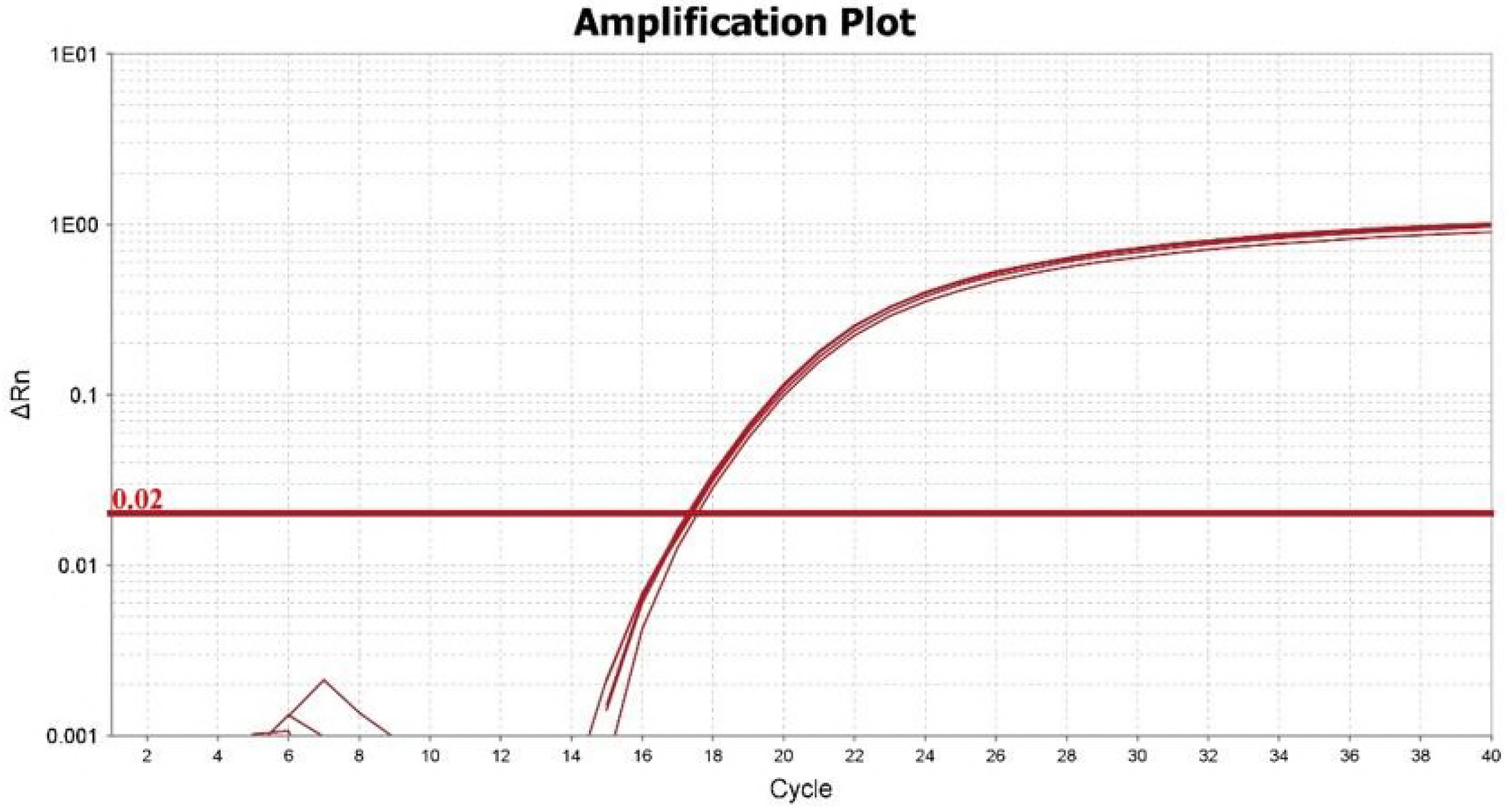
Specificity testing of the real-time qPCR assay using four *Serratia fonticola* strains.

### Standard curves and sensitivity of single real-time qPCR assays

Recombinant *invA-* and *gyrB*-containing plasmids were 10-fold serially diluted nine times, resulting in real-time qPCR amplicon copy numbers from 1. 97×10^10^ copies/μL to 1. 97×10^2^ copies/μL, and 1.45×10^10^ copies/μL to 1.45×10^2^ copies/μL, respectively. The standard curve for *invA* has a γ intercept of 48.116, a slope of −3.267, and a mean efficiency of 102.344% (Figure 3). The *gyrB* standard curve has a γ intercept of 42.919, a slope of −3.242, and a mean efficiency of 103.429% (Figure 4). As shown in Fig. 5 and Fig. 6, the single quantitative real-time PCR assays could detect *Salmonella* spp. and *S. fonticola* at concentrations as low as 197 and 145 copies per reaction, respectively.

**FIGURE 3.**
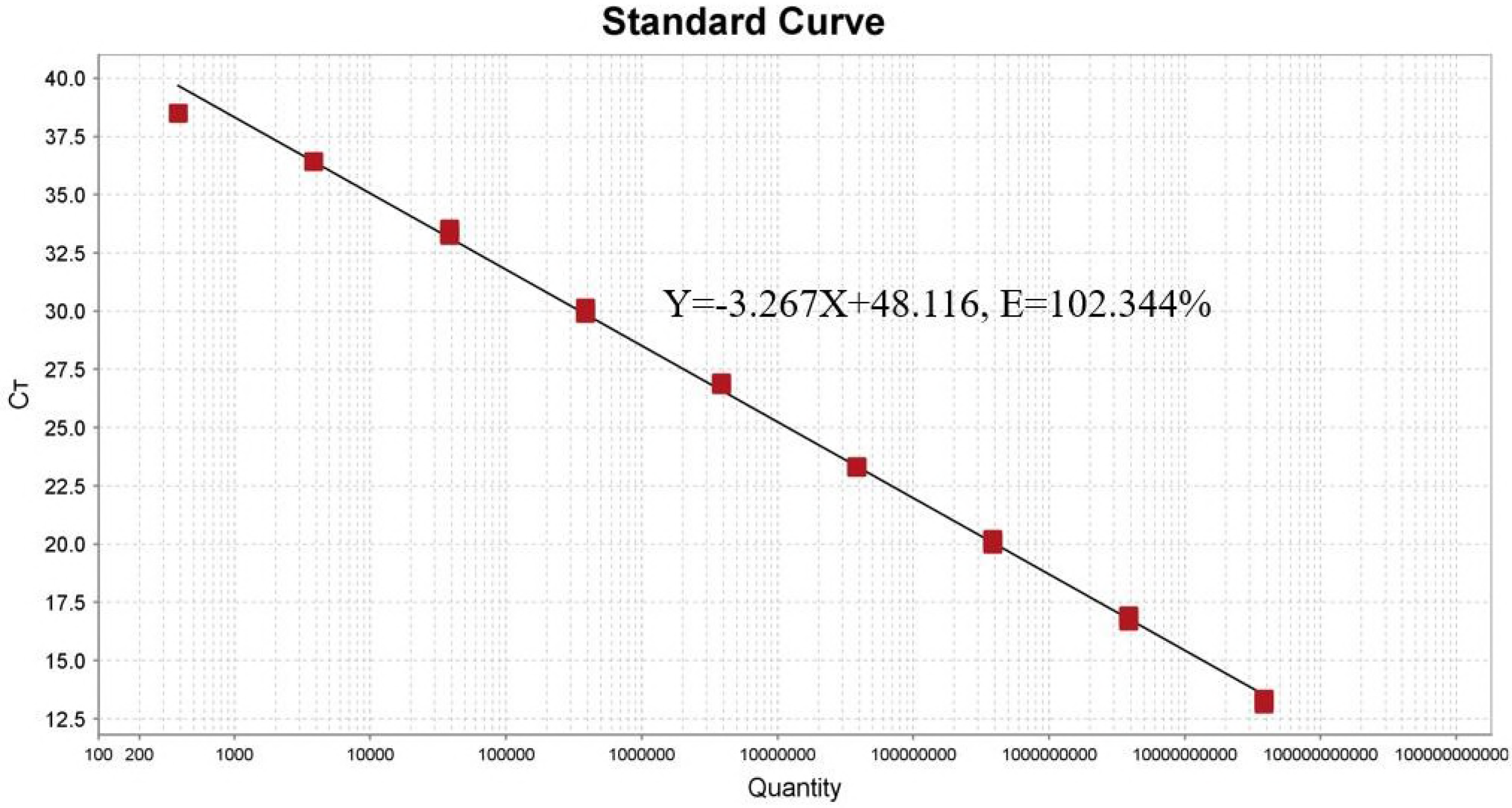
Standard curves for *Salmonella* spp. real-time qPCR assay. The curves represent a range from 10^10^ to 10^2^ copies per reaction.

**FIGURE 4.**
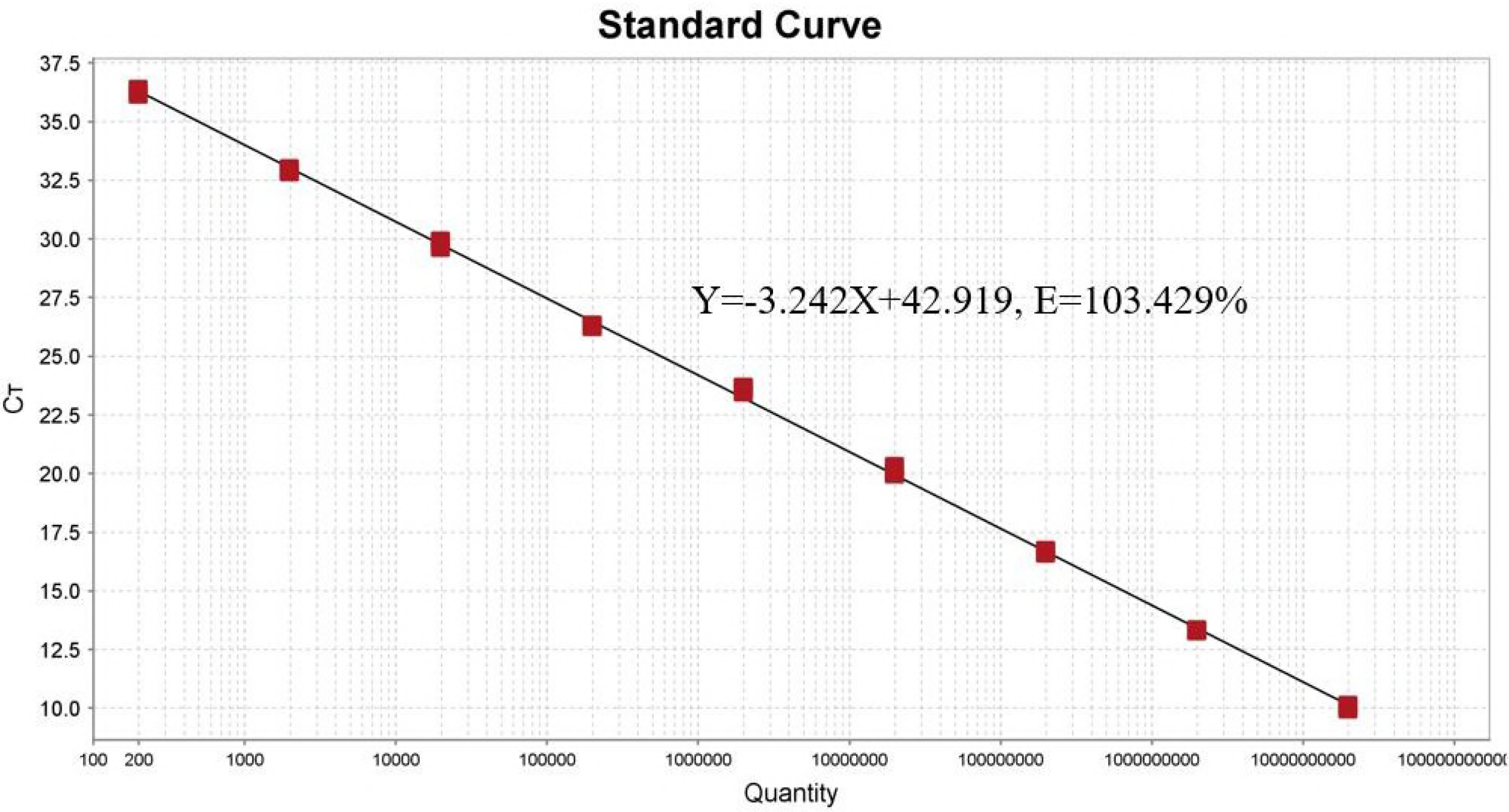
Standard curves for *S. fonticola* real-time qPCR assay. The curves represent a range from 10^10^ to 10^2^ copies per reaction.

**FIGURE 5.**
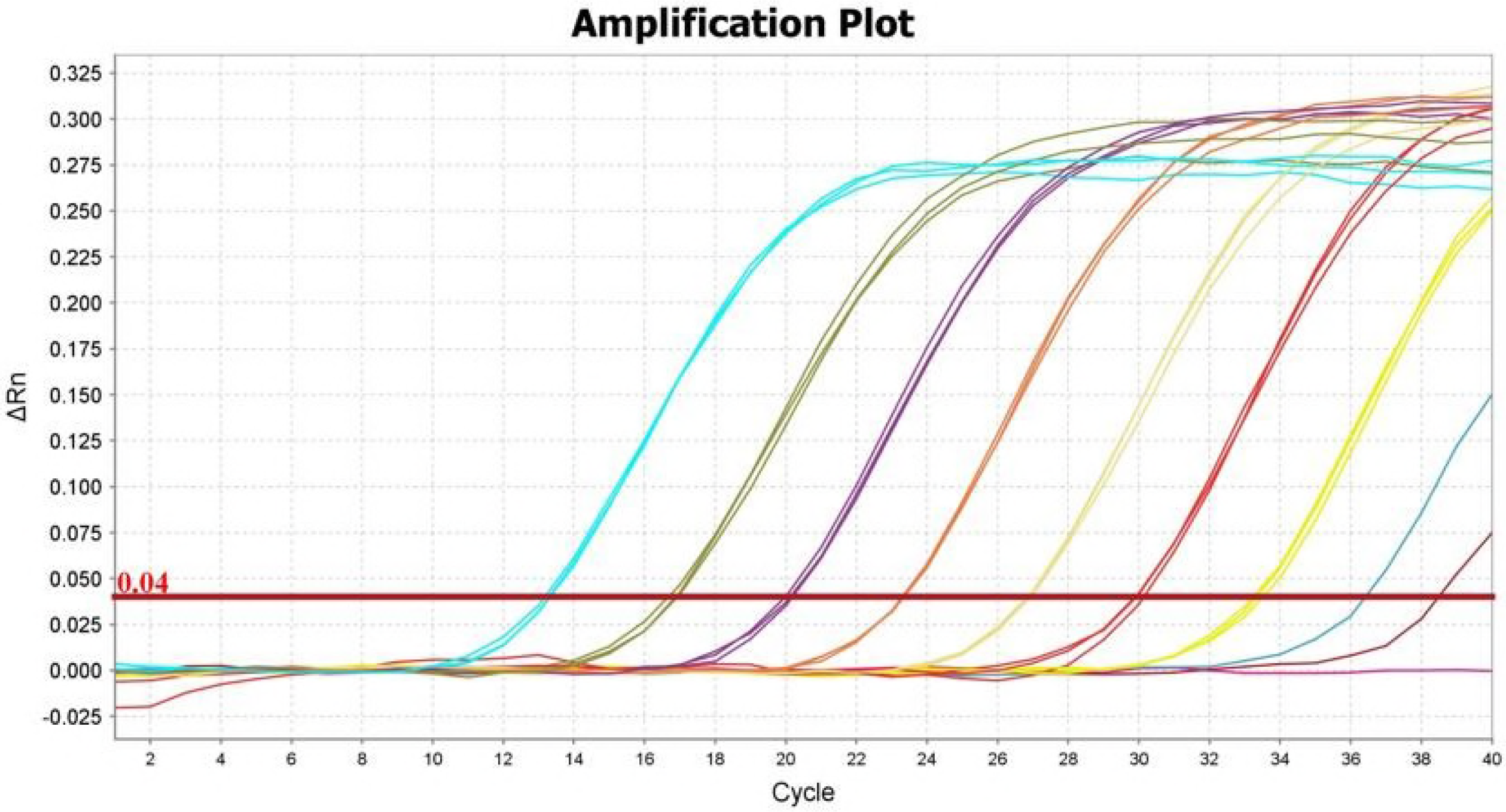
Sensitivity of the real-time qPCR assay for the detection of *Salmonella* species. Amplification plots from left to right represent a range of *invA* gene-containing plasmids from 1. 97×10^10^ to 1. 97×10^2^ copies/μL, respectively. Amplification plot 10 is a negative control.

**FIGURE 6.**
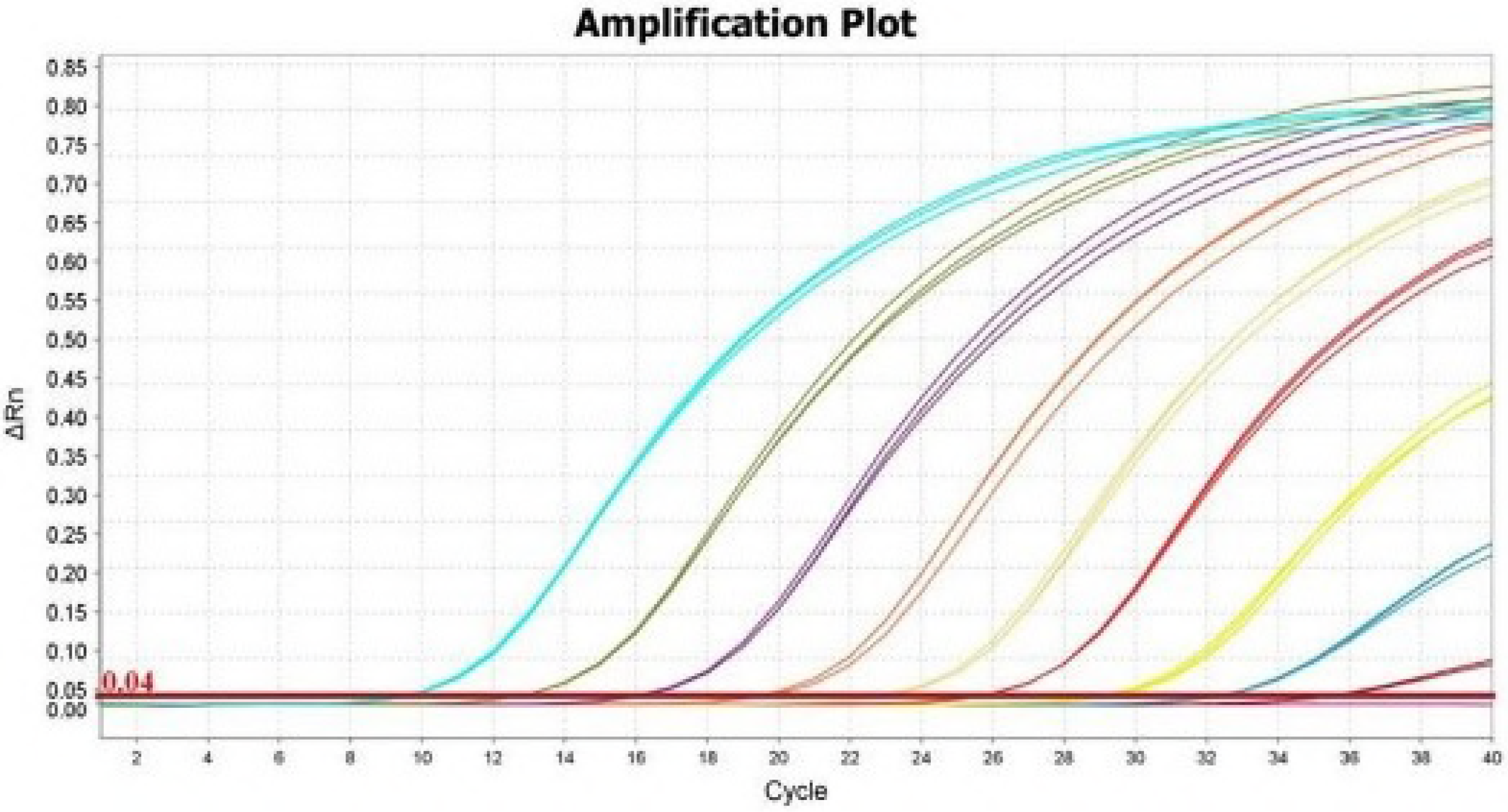
Sensitivity of the real-time qPCR assay for the detection of *S. fonticola.* Amplification plots from left to right represent a range of *gyrB* gene-containing plasmid from 1.45×10^10^ to 1.45×10^2^ copies/μL, respectively. Amplification plot 10 is a negative control.

### Establishment of the duplex real-time qPCR assay

Figure 7 and Figure 8 shows amplification plots and standard curves for the duplex real-time qPCR assay established for simultaneous detection of *Salmonella spp.* and *S. fonticola* developed with recombinant pUCm-*invA* and pUCm-*gyrB* plasmids. Standard curve slopes are −3.362 and −3.409 for the detection of *invA* and *gyrB,* respectively, indicating an amplification efficiency of 98.346% and 96.49%. A correlation coefficient consistently higher than 0.999 indicates effective simultaneous detection of two kinds of pathogens in one real-time qPCR assay without cross-reaction(data shown in Table5).

**TABLE 5.** Comparison of parameters between duplex and single real-time PCR assays

| Plasmid | Slope |  | y-axis intercept |  | R<sup>2</sup> |  | Efficiency ( % ) |  |
| --- | --- | --- | --- | --- | --- | --- | --- | --- |
|  | Single | Duplex | Single | Duplex | Single | Duplex | Single | Duplex |
| pUCm-*invA* | -3.267 | -3.362 | 48.116 | 48.775 | 0.998 | 0.999 | 102.344 | 98.346 |
| pUCm-*gyrB* | -3.242 | -3.409 | 42.919 | 44.396 | 0.999 | 0.999 | 103.429 | 96.49 |

**FIGURE 7.**
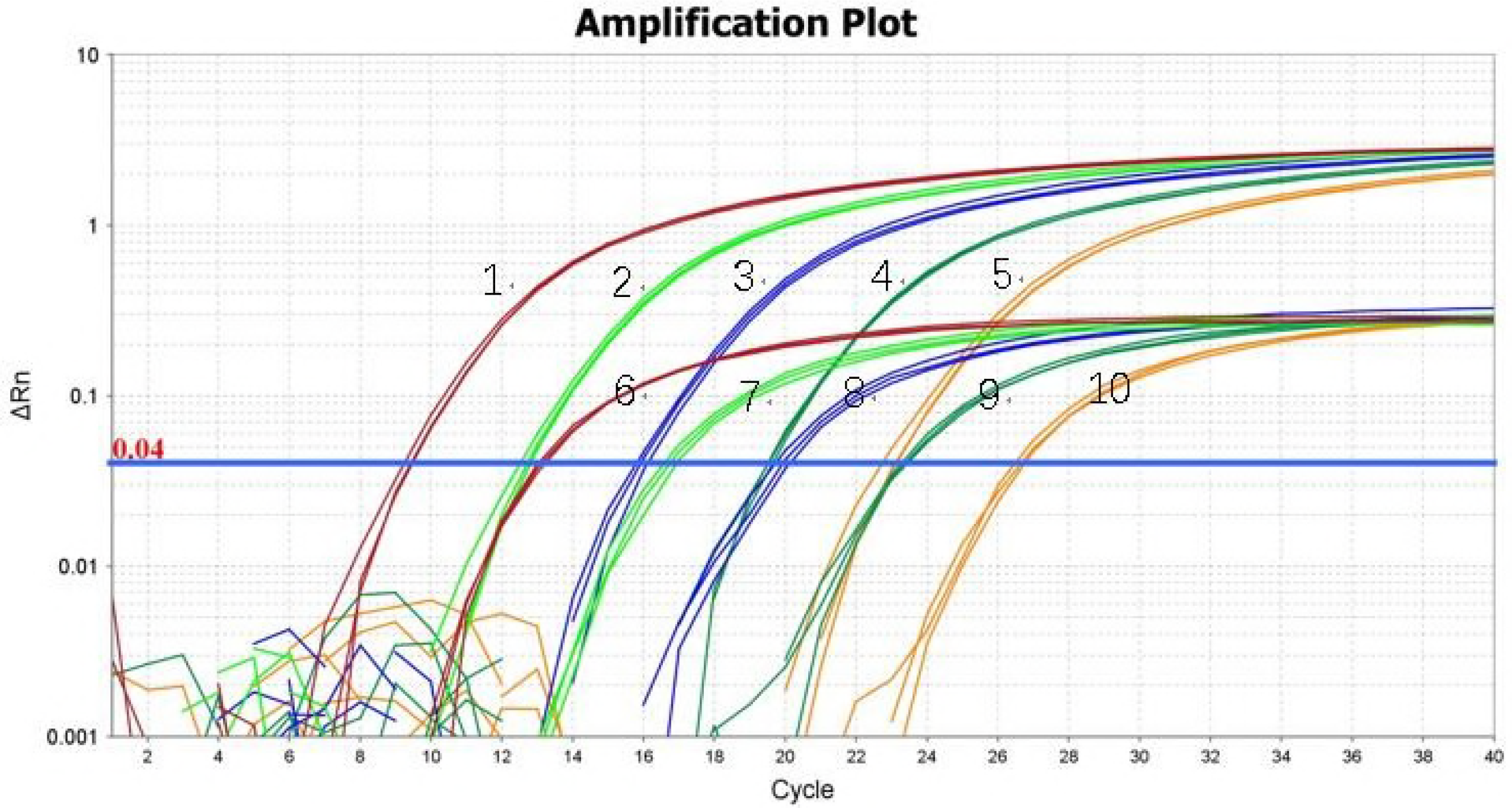
Amplification plot for the duplex real-time qPCR assay using serially diluted plasmids. Amplification plots 1 −5 represent *gyrB* gene-containing plasmid ranging from 1. 97×10^10^ to 1. 97×10^6^ copies/μL, respectively. Amplification plots 6-10 represent *invA* gene-containing plasmid ranging from 1. 45×10^10^ to 1. 45×10^6^ copies/μL, respectively.

**FIGURE 8.**
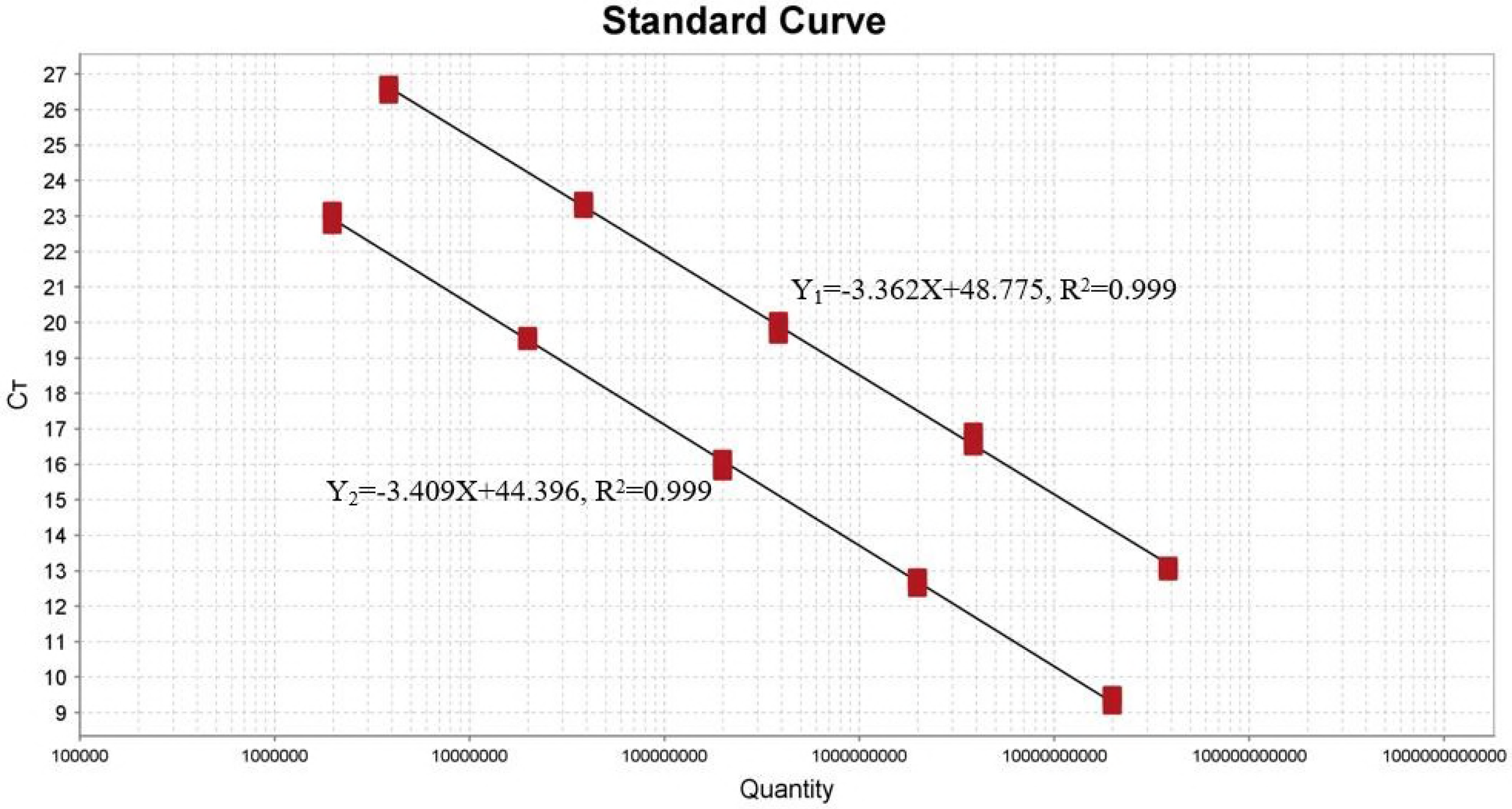
Standard curves for the duplex RT-qPCR assay. Y_1_ and Y_2_ are standard curves using *invA* gene- and *gyrB* gene-containing plasmids, respectively.

### Testing of reproducibility and stability

Five recombinant *invA* and *gyrB* plasmid dilutions were simultaneously used as substrates to evaluate the reproducibility and stability of the assay. As shown in Table 6, the standard deviation was no more than 0.153 for all reactions, and the coefficient of variation was less than 0.73%. Additionally, Ct values for the same plasmid samples obtained from same experiments were determined to analyze the performance of the developed real-time qPCR method. The results showed that amplification plots for all 96 replicate reactions were almost coincident in the vicinity of the threshold line (data not shown). Thus, the established quantitative real-time PCR assay has high stability and reproducibility.

**TABLE 6.** Reproducibility and stability testing of the duplex real-time PCR

| Ten-fold dilution<br>(copies/μL) | C<sub>t</sub> value | | | C<sub>t</sub> MN ± SD | CV (%) |
| --- | --- | --- | --- | --- | --- |
|  | 1 | 2 | 3 |  |  |
| 1.97E+08 | 19.989 | 20.086 | 20.195 | 20.090 ± 0.103 | 0.51 |
| 1.97E+07 | 23.287 | 23.331 | 23.302 | 23.307 ± 0.022 | 0.09 |
| 1.97E+06 | 26.924 | 26.861 | 26.879 | 26.879 ± 0.040 | 0.15 |
| 1.97E+05 | 29.892 | 29.934 | 29.910 | 29.910 ± 0.022 | 0.07 |
| 1.97E+04 | 33.251 | 33.525 | 33.356 | 33.377 ± 0.139 | 0.42 |
| 1.45E+08 | 15.997 | 16.005 | 15.856 | 15.952 ± 0.084 | 0.53 |
| 1.45E+07 | 19.581 | 19.330 | 19.343 | 19.418 ± 0.141 | 0.73 |
| 1.45E+06 | 22.909 | 22.724 | 22.898 | 22.844 ± 0.104 | 0.46 |
| 1.45E+05 | 25.522 | 25.526 | 25.628 | 25.559 ± 0.060 | 0.23 |
| 1.45E+04 | 28.918 | 29.209 | 29.146 | 29.091 ± 0.153 | 0.53 |

SD, Standard Deviation; MN, Mean; CV (Coefficient of Variance) = (SD / MN) × 100%

### Duplex real-time qPCR analysis of imported feedstuffs

Imported feedstuffs not infected with either *Salmonella* spp. or *S. fonticola* were artificially contaminated with the corresponding pathogens. Half of the feed samples were contained with *Salmonella* spp., while the other half were contained with *S. fonticola,* and genomic DNA was extracted for analysis after treatment. Detectable fluorescent signals were observed for all artificially contaminated feed samples following amplification. Furthermore, analysis of clinical imported fishmeal samples resulted in 10 amplification plots for the detection of *Salmonella* spp. and four amplification plots for the detection of *S. fonticola* (data not shown). Finally, the results of the detection of artificial and clinical samples were consistent with the national standard method (GB/T13091-2002) that was also employed for verification.

## Discussion

Many target genes for the detection of genus *Salmonella* species have been reported, including *afgA, hilA, spvC, sef* (33, 34), *fliC, fliB, iroB, rfbJ* (35), *ompC* (36), *spvR* (37), *fimA* (38), and *viaB* (39). However, they have shortcomings for the identification of *Salmonella* species, and false-positives limit the diagnostic process. At present, quorum sensing-related such as *luxS* and *gyrB* are widely used, along with virulence genes *invH, sopE, hilA,* and *invA* in the SP11 pathogenicity island, *sugR*, *rhuM*, and *iacP* in the SPI-3 pathogenicity island, and *spvB* and *spvC* in virulence plasmids. Of these, *invA,* which encodes an epithelial cell surface protein, is present in all *Salmonella* species, and is the most widely reported target gene in the *Salmonella* genus. Primers and probes for the diagnosis of *Salmonella* spp. based on this conserved gene are highly specific, hence *invA* was selected as the target gene for real-time qPCR detection of *Salmonella* species in the present study.

The 16S rDNA gene has been used for classification and identification of *Serratia* species in previous studies, as has the quorum sensing gene *luxS* (17), which was employed in real-time qPCR (21). In addition, several studies used carbapenem antibiotic resistance genes for identification of *Serratia* species (40). The *gyrB* gene, encoding the B subunit of DNA gyrase (GyrB) that forms topoisomerase II, an essential protein in replication, transcription, DNA synthesis, and maintenance of the DNA supercoiled structure, is more reliable than the 16S rDNA gene for the identification of bacterial species (41). The protein-coding *gyrB* gene contains more genetic information than the non-protein-coding 16S rDNA, resulting in a greater capacity to distinguish bacterial species (42). Thus, the *gyrB* gene was selected for discriminating *S. fonticola* and *Salmonella* spp. using real-time qPCR.

Real-time qPCR requires stricter primers, amplification products, and reaction conditions than conventional PCR. Primers used for real-time qPCR are typically 18-30 bp in length. Shorter primers (<15 nucleotides) can be combined efficiently, but often to the detriment of specificity. By contrast, longer primers display enhanced specificity but may also hybridize with the wrong pairing sequence, reducing specificity and decreasing the efficiency of hybridization, resulting in diminished PCR amplification. Ideal results are generally obtained when amplifying products of less than 300 bp. Therefore, the PCR products selected for differentiating *Salmonella* spp. and *S. fonticola* were 199 bp and 94 bp, respectively. Different annealing temperatures were tested to determine the optimum annealing temperature of the two pairs of primers, in order to reduce the influence of annealing temperature on the duplex real-time qPCR experiment. An annealing temperature of 64°C was found to be optimal.

The duplex real-time qPCR approach established for the rapid detection of *Salmonella* spp. and *S. fonticola* in imported feedstuffs was characterized by a high correlation coefficient between the Ct value and the logarithm of the initial copy number (R^2^ = 0. 999). Additionally, parameters including slope, y-axis intercept, and amplification efficiency between duplex real-time PCR and single real-time PCR were compared. As shown in Table 5, the slope of the standard curve and its intercept on the y-axis are approximately equal, and the amplification efficiency is ~100%. Thus, simultaneous detection of the two pathogens was achieved in a duplex real-time qPCR amplification assay, and the detection limit of this method is suitable for daily inspection and quarantine work. In summary, the rapid diagnostic method established in this study has many advantages, including a low detection limit and high repeatability. It is also rapid and convenient to deploy, since the results can be obtained within several hours after pre-enriching, representing a significant improvement in efficiency for detecting *Salmonella* spp. in imported animal-derived feedstuffs during quarantine work. This method is of great theoretical and practical value for ensuring the safety of imported feedstuffs, and could help to prevent adventive pathogens by facilitating effective quarantining and disposal measures.

## Authors and Contributors

Conceived and designed the experiments: Jh R, Wj W, and Dj Y Performed the experiments: Jh R. Generated and analyzed the data: Jh R, Wj W, Qy B, TZ, and Zd Z. Collected the clinical isolates: Ty Z, Sy Y, and Sj L. Wrote the paper: Jh R. Revised the paper: Dj Y and Bx Z. All authors read and approved the final manuscript.

## Acknowledgements

We would like to thank all the teachers at Fujian Entry-exit Inspection and Quarantine Bureau, Fujian Key Laboratory for Technology Research of Inspection and Quarantine in Fujian Province, China. And the native English speaking scientists of Elixigen Company (Huntington Beach, California) for editing our manuscript.

